# Ridge regression baseline model outperforms deep learning method for cancer genetic dependency prediction

**DOI:** 10.1101/2023.11.29.569083

**Authors:** Daniel Chang, Xiang Zhang

## Abstract

Accurately predicting genetic or other cellular vulnerabilities of unscreened, or difficult to screen, cancer samples will allow vast advancements in precision oncology. We re-analyzed a recently published deep learning method for predicting cancer genetic dependencies from their omics profiles. After implementing a ridge regression baseline model with an alternative, simplified problem setup, we achieved a model that outperforms the original deep learning method. Our study demonstrates the importance of problem formulation in machine learning applications and underscores the need for rigorous comparisons with baseline approaches.

## Main

Precision oncology methods rely on the ability to accurately translate molecular measurements of tumors to insights about their genetic dependencies or other cellular vulnerabilities, ultimately dictating targeted therapeutics. Through genome-wide CRISPR-Cas9 knockout screens, the Cancer Dependency Map(*1*–*5*) (DepMap) has characterized the genetic dependency profiles of over 1000 cancer cell lines (CCLs). Using the DepMap 2018Q2 release, Chiu *et al*. recently developed DeepDEP, a deep learning method for predicting genetic dependency profiles of CCLs given their multi-omics(*6*). DeepDEP was reported to vastly outperform baseline conventional machine learning models. However, we argue here that this result can be attributed to aspects of its problem formulation.

Notably, DeepDEP does not jointly predict the entire genetic dependency profile of an input CCL at once. Instead, the model takes as input both multi-omics of a CCL and a functional fingerprint of a single gene dependency of interest (DepOI, as abbreviated by Chiu *et al*.), and outputs the predicted score of that specific gene DepOI for that CCL (Fig. 1A). Functional fingerprints were defined as binary vectors encoding a gene’s involvement in 3115 chemical and genetic perturbations (CGPs) from MSigDB v6.2(*7*), potentially facilitating the model to learn relationships between genes with functional similarities.

**Fig. 1.**
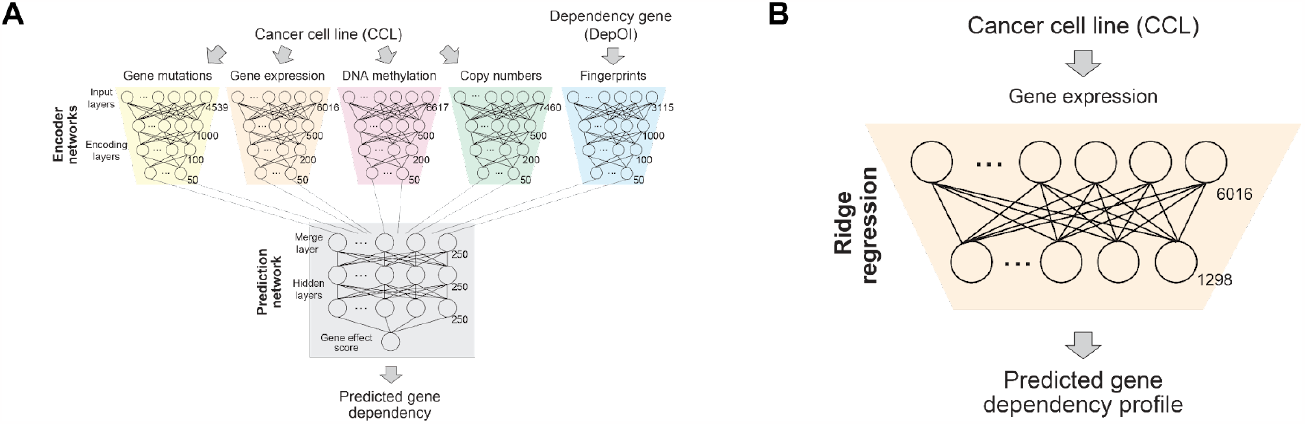
Comparison of DeepDEP and baseline setups. **(A)** DeepDEP architecture (taken directly from Chiu *et al*.). DeepDEP takes as input a CCL’s DNA mutation, gene expression, DNA methylation, and copy number alteration profiles. In addition, a functional fingerprint of a gene dependency of interest (DepOI) is supplied. Dimensionality of each input is displayed. DeepDEP performs dimensionality reduction using an autoencoder pretrained on 8238 TCGA tumors. The model then merges the dimensionality-reduced data into a prediction network to predict the score of the gene DepOI (corresponding to the input functional fingerprint) for a given CCL. **(B)** Baseline model setup. A multioutput ridge regression model takes as input omics data (in this illustration, just gene expression) from a CCL, and predicts its genetic dependency profile across 1298 cancer relevant gene DepOIs, as defined by Chiu *et al*.

We recognized that this problem formulation, while likely beneficial for embedding prior knowledge about a gene DepOI’s function in a deep learning context, could be problematic for simpler baseline models as it requires a model to learn highly non-linear relationships between input omics features and dependency scores. Because this formulation requires a singular model to predict the score of any gene DepOI (given its functional fingerprint representation), the model is unable to directly relate input omics features to dependency scores. Instead, an optimal model must first generate intermediate representations composed of both a CCL’s input omics features and the DepOI’s fingerprint vector. Because Chiu *et al*. evaluated baseline model performances using this setup, we were skeptical about the degree by which DeepDEP truly outperforms baseline methods.

To evaluate this, we implemented a ridge regression baseline model using a simplified problem formulation (Fig. 1B). Instead of outputting a single gene dependency score for a CCL and gene DepOI combination, our baseline jointly outputs predictions of an input CCL’s genetic dependency profile across 1298 cancer relevant genes (defined by Chiu *et al*.), the same gene DepOI set on which DeepDEP was trained and evaluated.

We compared 10-fold cross validation results of DeepDEP and a ridge regression baseline using the aforementioned simplified setup. The ridge regression baseline here uses all input omics features, but no functional fingerprints. Across 278 CCLs and 1298 gene DepOIs, DeepDEP and the ridge regression baseline achieved similar predictive performances (DeepDEP: Fig. 2A, Pearson correlation coefficient ρ = 0.87; ridge regression baseline: Fig. 2B, ρ = 0.88). However, because of the existence of pan-essential genes, for which dependency score predictions are mostly constant across CCLs and thus easy to predict, analyzing correlation results across all genes likely yields an overly-optimistic estimate of model performances. Thus, we next evaluated the results by computing correlations per-gene-DepOI. DeepDEP achieved a mean per-gene-DepOI ρ of 0.137 (Fig. 2C), while the ridge regression baseline achieved a ρ of 0.276 (Fig. 2E). The ridge regression baseline (with this simplified setup) not only achieved a result higher than that of any baseline machine learning methods evaluated by Chiu *et al*. (not shown here), but also outperformed the full DeepDEP model.

**Fig. 2.**
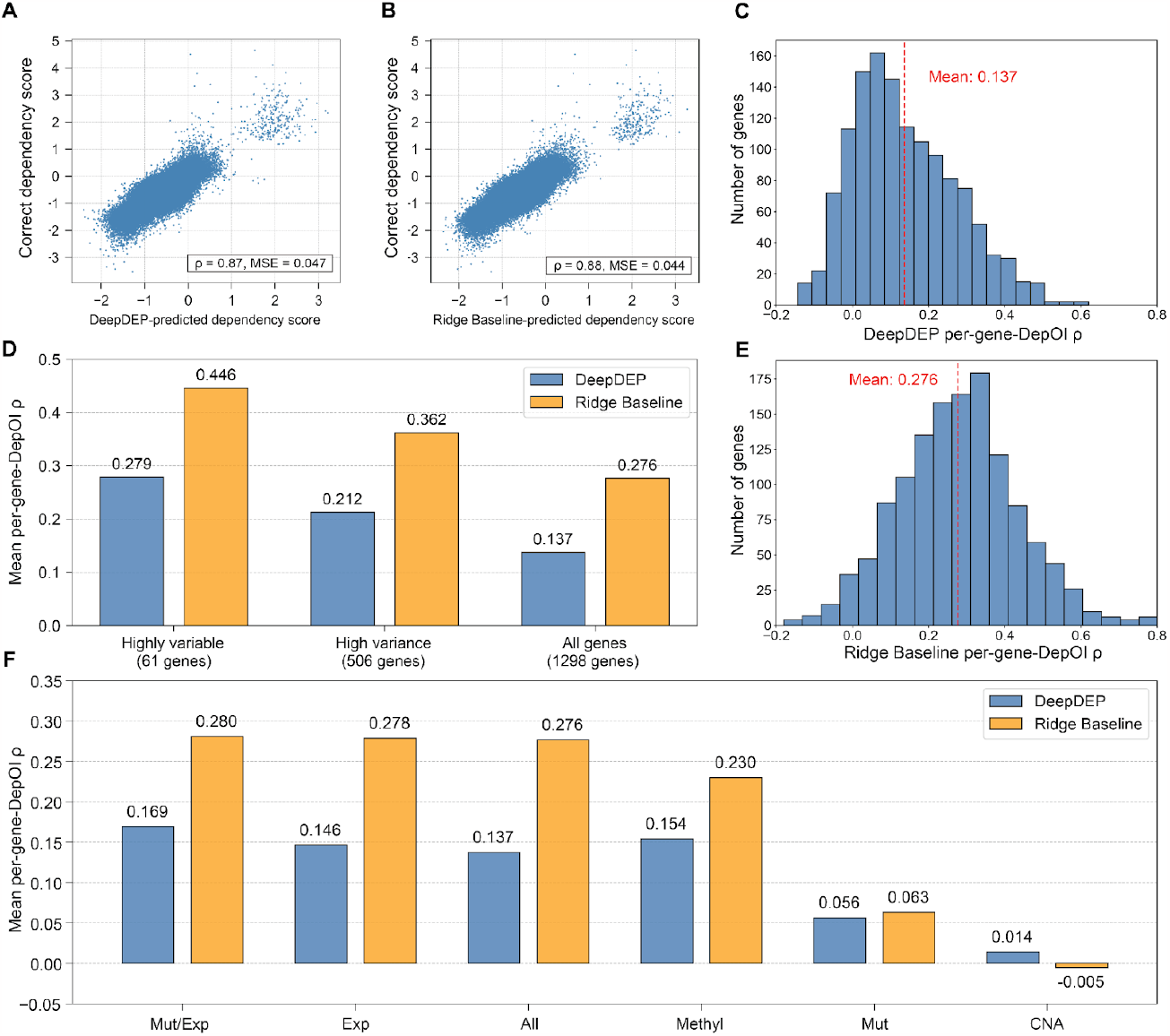
Ridge regression baseline outperforms DeepDEP. **(A** and **B)** Scatterplots of DeepDEP (x-axis, **A**) and ridge baseline (x-axis, **B**) predicted dependency scores vs the correct dependency scores (y-axis) across 278 CCLs and 1298 gene DepOIs. 10-fold cross-validation, where CCLs are held out, was used to generate predictions for both models. DeepDEP achieves ρ = 0.87 and mean squared error (MSE) = 0.047. The ridge baseline achieves ρ = 0.88 and MSE = 0.044. (**C** and **E**) Histogram of per-gene-DepOI ρ for DeepDEP and the ridge baseline. DeepDEP achieves a mean ρ of 0.137, and the ridge baseline achieves a mean ρ of 0.276. **(D)** Mean per-gene-DepOI ρ for (as defined in Chiu *et al*.) highly variable dependency score genes (n = 61), high variance dependency score genes (n = 506), and the entire gene set (n = 1298). The ridge regression baseline model outperforms DeepDEP on all gene sets. **(F)** Simplified models trained on subsets of the omics data. Mut = mutation, Exp = gene expression, Methyl = methylation, and CNA = copy number alteration. For all simplified models except that trained on just CNA data, the ridge regression baseline achieves a higher mean per-gene-DepOI ρ than DeepDEP does.

Chiu *et al*. also analyzed mean per-gene-DepOI ρ on two subsets of genes that were observed to have high variance dependency scores and thus were likely cancer-relevant genes. The ridge regression baseline model outperformed DeepDEP on both of these gene sets (Fig. 2D).

We next constructed several simplified ridge models using subsets of the omics data types (for example, Fig. 1B depicts an expression-only ridge model), similarly as done in Chiu *et al*. These simplified models were compared with DeepDEP results. In all instances except when using only copy number alteration data, the ridge regression baseline achieves a higher mean per-gene-DepOI ρ than DeepDEP does (Fig. 2F).

These results demonstrate how machine learning problem formulations dramatically impact the performance of baseline approaches. We show that a problem formulation that is convenient for embedding information into deep learning models may not always be the ideal formulation for baseline approaches. To truly evaluate the degree by which novel methods outperform baselines, it is necessary to rigorously evaluate baselines using different, potentially simpler, problem formulations. Moreover, many of our simplified ridge regression baselines, trained on subsets of the data types (Mut/Exp, Exp, Methyl), outperformed DeepDEP trained on all input genomic features. This demonstrates that the proposed deep learning model was not able to benefit from integrating information from a diverse set of data modalities. We also demonstrate how the use of gene functional fingerprints is not important for achieving an elevated prediction performance over DeepDEP. These results bring into question the strength of the results obtained from downstream analyses using the DeepDEP model, such as the pan-cancer tumor dependency map that Chiu *et al*. generated by applying DeepDEP on TCGA data.

Overall, we demonstrate the importance of conducting rigorous baselines when evaluating the performance of novel methods, and the importance of considering the implications of different machine learning problem formulations.

## Data and code availability

All data was obtained from the Code Ocean compute capsule accompanying Chiu *et al*., the supplementary tables of Chiu *et al*., and from the DepMap portal. Machine learning tasks were performed using Scikit-learn v1.2.2(*8*). All code and data for this analysis can be found at: https://github.com/danielchang2002/deepdep_reanalysis.

## Methods

DeepDEP CCL dependency score predictions from 10-fold cross-validation (where CCLs are held out) were obtained from Supplementary Table S8 of Chiu *et al*. Ground truth gene effect scores of the 278 CCLs were obtained from the DepMap 2018Q2 release. The DeepDEP 10-fold cross-validation dependency score predictions were compared with the ground truth in Fig. 2A and 1C.

DNA mutation, gene expression, DNA methylation, and copy number alteration data of the 278 CCLs were obtained from the Code Ocean compute capsule accompanying Chiu *et al*. Functional fingerprints were not used in our analysis. Ridge regression (sklearn.linear_model.Ridge) with default parameters was then used to predict an input CCL’s genetic dependency profile across 1298 cancer relevant genes (defined by Chiu *et al*.), the same gene DepOI set on which DeepDEP was trained and evaluated, given a flattened vector of the four data types concatenated together for an input CCL. Ridge regression baseline prediction scores were obtained via 10-fold cross-validation (where CCLs are held out). Notably, the CCL fold partitioning is not identical to that used to generate DeepDEP cross-validation results in Supplementary Table S8 of Chiu *et al*., but generated independently in this analysis (sklearn.model_selection.KFold; random_state = 42). Ridge regression baseline 10-fold cross-validation scores were compared with the ground truth in Fig. 2B and 2E.

The gene set of 61 “highly variable” genes was obtained by locating genes with dependency score standard deviations greater than 0.3, as defined by Chiu *et al*. The gene set of 506 “high variance” genes was obtained using the “Achilles_high_variance_genes.csv” file provided by the DepMap portal. The 10-fold cross-validation results on these two gene subsets, for both the Ridge regression baseline and DeepDEP, are detailed in Fig. 2D.

Results for simplified ridge regression baseline models were obtained using subsets of the CCL omics data. This was performed identically as before, using 10-fold cross-validation. However, for DeepDEP, the 10-fold cross-validation prediction scores were only available (in the supplementary of Chiu *et al*.) for the model using all omics features (i.e. the “All” model), and not for the simplified models. Thus, in Fig. 2F, simplified DeepDEP model performances were obtained using data from Supplementary Fig. S5 of Chiu *et al*., which details per-gene-DepOI ρ for simplified models using 10 independent train-test subsampling (i.e., a slightly different evaluation method than 10-fold cross-validation).

